# Targeting redox regulatory site of protein kinase B impedes neutrophilic inflammation in lung injury

**DOI:** 10.1101/264184

**Authors:** Po-Jen Chen, I-Ling Ko, Chia-Lin Lee, Hao-Chun Hu, Fang-Rong Chang, Yang-Chang Wu, Yann-Lii Leu, Chih-Ching Wu, Cheng-Yu Lin, Chang-Yu Pan, Yung-Fong Tsai, Tsong-Long Hwang

**Author notes:** **Correspondence to:** Dr. Tsong-Long Hwang, Graduate Institute of Natural Products, Chang Gung University, 259 Wen-Hwa 1^st^ Road, Kweishan, Taoyuan 333, Taiwan.

## Abstract

Neutrophil activation has a pathogenic effect in inflammatory diseases. Protein kinase B (PKB)/AKT regulates diverse cellular responses. However, the significance of AKT in neutrophilic inflammation is still not well understood. Here, we identified CLLV-1 as a novel AKT inhibitor. CLLV-1 inhibited respiratory burst, degranulation, chemotaxis, and AKT phosphorylation in activated human neutrophils and dHL-60 cells. Significantly, CLLV-1 blocked AKT activity and covalently reacted with AKT Cys310 *in vitro*. The AKT_309-313_ peptide-CLLV-1 adducts were determined by NMR or mass spectrometry assay. The alkylation agent-conjugated AKT (reduced form) level was also inhibited by CLLV-1. Additionally, CLLV-1 ameliorated lipopolysaccharide (LPS)-induced acute lung injury (ALI) in mice. CLLV-1 acts as a covalent allosteric AKT inhibitor by targeting AKT Cys310 to restrain inflammatory responses in human neutrophils and LPS-induced ALI *in vivo*. Our findings provide a mechanistic framework for redox modification of AKT that may serve as a novel pharmacological target to alleviate neutrophilic inflammation.

## Introduction

Neutrophils are the first line of host defense in the innate immune response. They are chemoattracted to inflammatory regions in response to infection, and they subsequently eliminate invading pathogens through respiratory burst, degranulation, and neutrophil extracellular traps (NETs). However, overwhelming neutrophil activation plays a critical role both in infective and sterile inflammation *(Jorch & Kubes, 2017; Leiding, 2017; Nauseef & Borregaard, 2014*). The reactive oxygen species (ROS) and proteases released by activated neutrophils can damage healthy surrounding tissues, resulting in deleterious inflammatory diseases, such as acute respiratory distress syndrome (ARDS), chronic obstructive pulmonary disease (COPD), sepsis, or asthma *(Aschner et al, 2014; Delano & Ward, 2016; Soehnlein et al, 2017*).

Pathogen recognition or an inflammatory environment triggers many critical intracellular signal processes through surface receptors in neutrophils *(Liu et al, 2016; Tsai et al, 2016; Yang & Hwang, 2016*). The serine/threonine-specific protein kinase, protein kinase B (PKB)/AKT, has been reported to regulate the neutrophil immune responses, including respiratory burst, degranulation, and chemotaxis *(Chamcheu et al, 2017; Manning & Toker, 2017; Zhang et al, 2013*). In human neutrophils, activated AKT phosphorylates p47^phox^, a component of nicotinamide adenosine dinucleotide phosphate (NADPH) oxidase, to initiate respiratory burst *(Chen et al, 2010; El-Benna et al, 2009; Hoyal et al, 2003*). Pharmacological inhibition of phospho-phosphoinositide 3-kinase (PI3K)/AKT signaling reduces leukocyte degranulation (*Hoenderdos et al, 2016; Nanamori et al, 2007*). AKT also stabilizes F-actin polymerization to enhance the chemotaxis of activated neutrophils (*Chen et al, 2010; Chodniewicz & Zhelev, 2003; Kumar et al, 2014*). Therefore, AKT may be a potential pharmacological target to treat neutrophilic inflammation. It is activated via the phosphorylation of Thr308 and Ser473 residues, and it is conversely deactivated through dephosphorylation *(Brazil et al, 2004; Weichhart et al, 2015*). In addition to the well-known regulatory phosphorylation of AKT, emerging evidence has supported the significance of redox modification of AKT *(Corcoran & Cotter, 2013; Zeng et al, 2018*). Along with through dephosphorylation, AKT is inactivated through an intra-disulfide bond between Cys296 and Cys310 in the catalytic domain *(Ahmad et al, 2014; Durgadoss et al, 2012; Murata et al, 2003*). However, the mechanistic details of whether redox-controlled AKT activity contributes to neutrophilic inflammation remains to be explored.

In this study, we identified that 5,7-dimethoxy-1,4-phenanthrenequinone (CLLV-1) (Figure 1A) is a novel AKT inhibitor that acts via a thiol-based reaction with the Cys310 residue of AKT to block its kinase activity. Phenanthrenequinones have been shown to exhibit antiplatelet aggregation and anticancer activity (*Lee et al, 2012; Lee et al, 2014*). Here, we found that CLLV-1 has an anti-inflammatory potential to impede respiratory burst, degranulation, and chemotaxis in activated human neutrophils or neutrophil-like differentiated HL-60 (dHL-60) cells. Moreover, administration of CLLV-1 and MK-2206, an allosteric AKT inhibitor, attenuated the inflammatory responses of lipopolysaccharide (LPS)-induced acute lung injury (ALI) *in vivo*. Our findings elucidate that redox modification of AKT may be a novel pharmacological strategy for suppressing neutrophil-dominant disorders. We also suggest that CLLV-1 has the potential to be developed as an anti-inflammatory drug.

**Figure 1.**
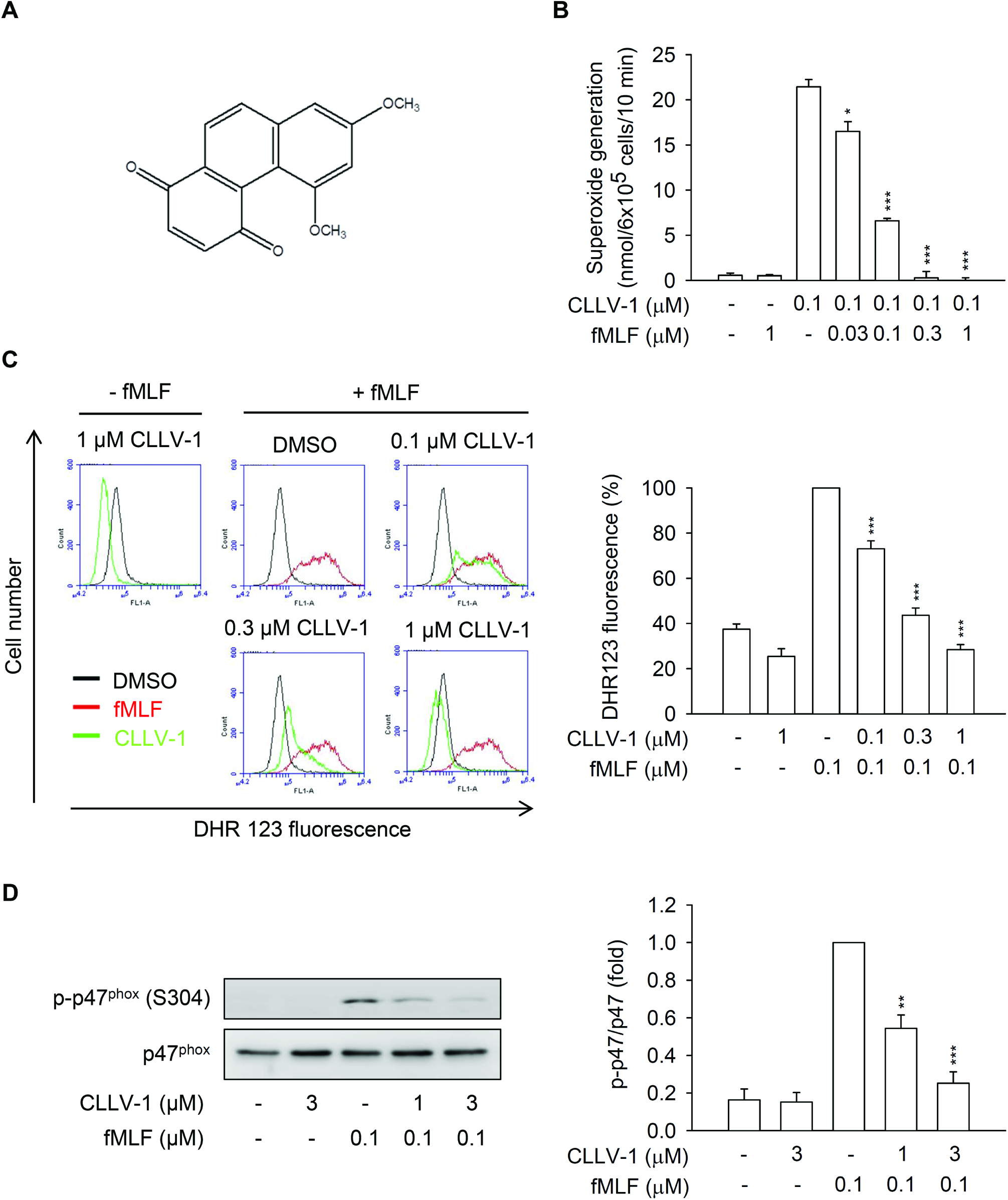
CLLV-1 attenuates superoxide anion generation, ROS formation, and p47^phox^ phosphorylation in fMLF-activated human neutrophils. (A) The chemical structure of CLLV-1. (B-C) Human neutrophils were preincubated with DMSO or CLLV-1 (0.03–3 μM) and then activated with or without fMLF (0.1 μM)/CB (1 μg/mL). (B) Superoxide anion generation was detected using cytochrome *c* reduction by a spectrophotometer at 550 nm. (C) The intracellular ROS was monitored by flow cytometry, using cell-permeable DHR123. (D) Phosphorylation of p47^phox^ was analyzed by immunoblotting, using antibodies against the phosphorylated (S304) and total p47^phox^. All data are expressed as mean values ± SEM (n = 3); \**p* < 0.05, \*\**p* < 0.01, and \*\*\**p* < 0.001 compared with the DMSO + fMLF group (Student’s *t*-test).

## Results

### CLLV-1 suppresses the inflammatory responses in fMLF-activated human neutrophils

To investigate the anti-inflammatory ability of CLLV-1, we first examined the effects of CLLV-1 on *N*-formyl-Met-Leu-Phe (fMLF)-induced respiratory burst, including superoxide anion production, ROS formation, and NADPH oxidase activation (p47^phox^ phosphorylation) in human neutrophils. CLLV-1 dose-dependently inhibited superoxide anion generation and ROS formation with IC_50_ values of 0.058 ± 0.006 and 0.106 ± 0.022 μM, respectively (Figure 1B and 1C). It did not induce LDH release, suggesting that it did not cause membrane damage and cytotoxicity. We further evaluated how CLLV-1 inhibited the superoxide anion generation in fMLF-activated human neutrophils. In a cell-free xanthine/xanthine oxidase system, CLLV-1 (0.1–3 μM) did not exhibit a superoxide anion-scavenging ability. Superoxide dismutase (SOD) was a positive control. Superoxide anion is produced by NADPH oxidase in human neutrophils (*El-Benna et al, 2009*). The isolated neutrophil membrane and cytosol fractions were used to examine the inhibitory effect of CLLV-1 on NADPH oxidase: CLLV-1 (0.3 and 3 μM) did not reduce superoxide anion production in SDS-reconstituted NADPH oxidase (Figure 1-figure supplement 1A-C). Diphenyleneiodonium (DPI; 10 μM), an NADPH oxidase inhibitor, was a positive control. The phosphorylation of p47^phox^, a component of NADPH oxidase, was repressed by CLLV-1 in fMLF-activated human neutrophils (Figure 1D), suggesting that the anti-inflammatory effect of CLLV-1 on respiratory burst may be through modulating upstream signaling of NADPH oxidase in human neutrophils.

Next, the effects of CLLV-1 on human neutrophil degranulation and chemotaxis were determined. CLLV-1 repressed fMLF-induced elastase release with an IC_50_ value of 0.172 ± 0.011 μM (Figure 2A). In contrast, it failed to alter the activity of elastase in a cell-free assay (Figure 2-figure supplement 1D), suggesting that it inhibited human neutrophil degranulation via the regulation of intracellular signaling pathways. In addition, integrin CD11b activation leads to neutrophils adhering to endothelial cells and subsequently induces neutrophil migration and infiltration. F-actin polymerization at the leading edge in polarized neutrophils governs the chemotaxis (*Chodniewicz & Zhelev, 2003*). CLLV-1 decreased CD11b expression and F-actin assembly in fMLF-activated human neutrophils and suppressed fMLF-induced neutrophil adhesion to bEnd.3 endothelial cells (ECs) and migration (Figure 2B-E-figure supplement 2).

**Figure 2.**
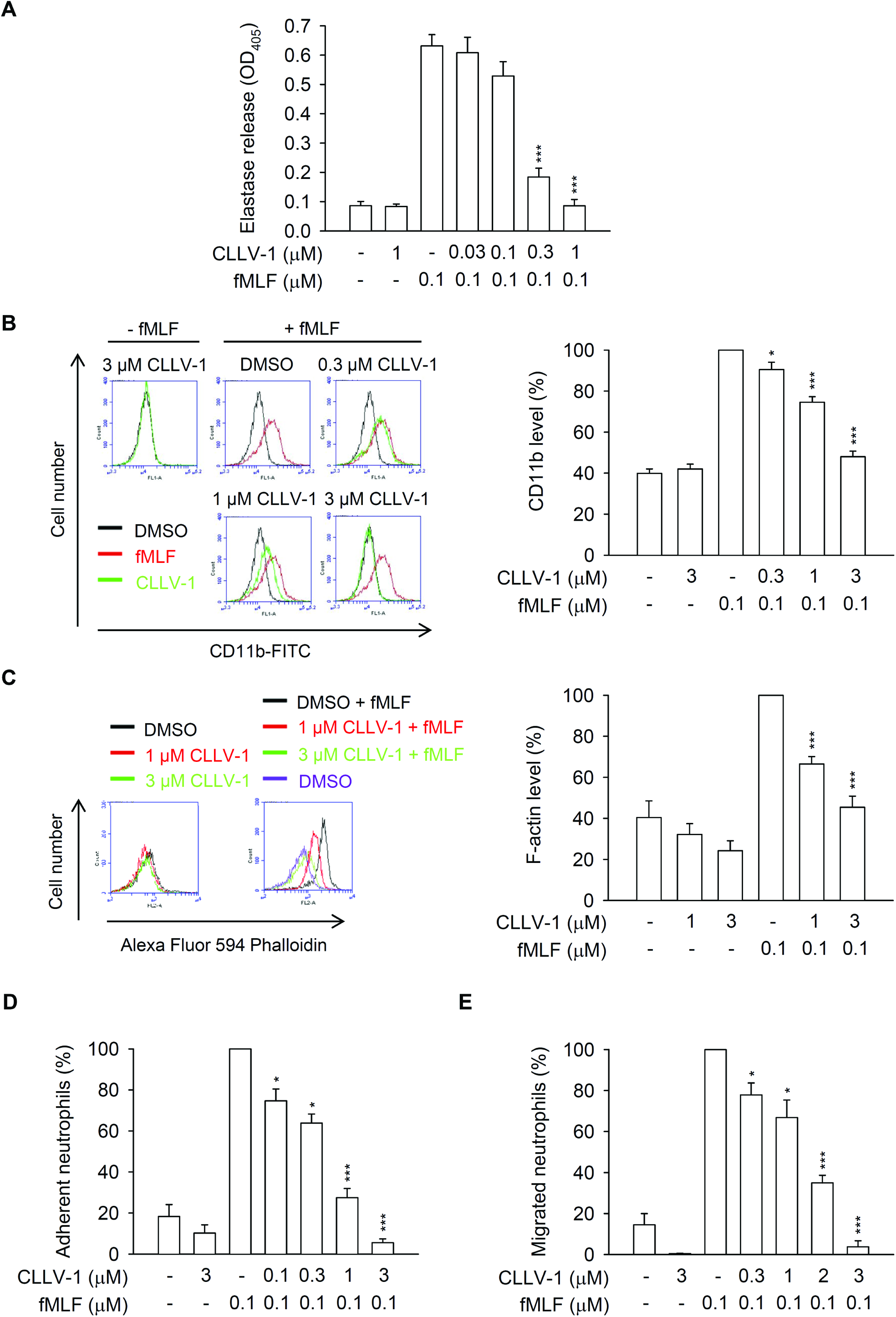
CLLV-1 inhibits elastase release, CD11b/F-actin expression, and neutrophil adhesion/migration in fMLF-activated human neutrophils. (A-C) Human neutrophils were incubated with DMSO or CLLV-1 (0.3–3 μM) for 5 min before stimulation with or without fMLF (0.1 μM)/CB (0.5 or 1 μg/mL). (A) Elastase release was measured by a spectrophotometer at 405 nm, using elastase substrate. (B) The CD11b levels on cell surface were detected by flow cytometry, using FITC-labeled anti-CD11b antibodies. (C) The F-actin levels were assayed by flow cytometry, using Alexa Fluor 594 Phalloidin. (D) Hoechst 33342-labeled neutrophils were pretreated with DMSO or CLLV-1 (0.1–3 μM) for 5 min and stimulated with fMLF (0.1 μM)/CB (1 μg/mL). Sequentially, neutrophils were incubated with LPS-preactivated ECs for another 30 min. After gently washing, EC-associated neutrophils were counted under a microscope. (E) Neutrophils were preincubated with DMSO or CLLV-1 (0.3–3 μM) for 5 min in the top chamber. Migrated neutrophils in the bottom wells with or without fMLF were counted after 90 min. All data are expressed as mean values ± SEM (n = 3); \**p* < 0.05, ^**^*p* < 0.01, and \*\*\**p* < 0.001 compared with the DMSO + fMLF group (Student’s *t*-test).

### CLLV-1 ameliorates AKT activation in response to various stimuli in human neutrophils

This study aimed to identify the target protein of CLLV-1 in human neutrophils. The fMLF mainly binds to formyl peptide receptor 1 (FPR1) to activate neutrophils through multiple intracellular signaling pathways such as AKT and mitogen-activated protein kinases (MAPKs) (*Dorward et al, 2015*). CLLV-1 (0.1–3 μM) did not compete with the fluorescently labeled fMLF analog *N*-formyl-Nle-Leu-Phe-Nle-Tyr-Lys (fNLFNYK) for FPR1 binding (Figure 3-figure supplement 1E), ruling out the effect of CLLV-1 on FPR1. Therefore, the activation of AKT, ERK, JNK, and p38 was examined in fMLF-activated human neutrophils. CLLV-1 inhibited the phosphorylation of AKT (Thr308 and Ser473), but not of ERK, JNK, or p38 (Figure 3). Because CLLV-1 selectively restrained AKT activation, we wondered whether CLLV-1 suppressed AKT activation and inflammatory responses in different stimuli-activated human neutrophils, including sodium fluoride (NaF; direct G protein activator), WKYMVm (FPR2 agonist), interleukin-8 (IL-8), and leukotriene B4 (LTB_4_) (*Futosi et al, 2013*). Further, CLLV-1 significantly inhibited NaF- and WKYMVm-induced superoxide anion generation in human neutrophils. It also showed an inhibitory effect on elastase release in NaF-, WKYMVm-, IL-8-, and LTB_4_-activated human neutrophils (Figure 3-figure supplement 3). Notably, it significantly suppressed the phosphorylation of AKT (Thr308 and Ser473) in human neutrophils activated by all tested stimuli (Figure 3-figure supplement 4), suggesting that AKT may be the target of CLLV-1 in human neutrophils. Another potent AKT inhibitor, MK-2206 (*Hirai et al, 2010*), also inhibited the superoxide anion generation and elastase release in fMLF-activated human neutrophils (Figure 3-figure supplement 5), supporting that inhibition of AKT is a potential strategy to attenuate neutrophilic inflammation.

**Figure 3.**
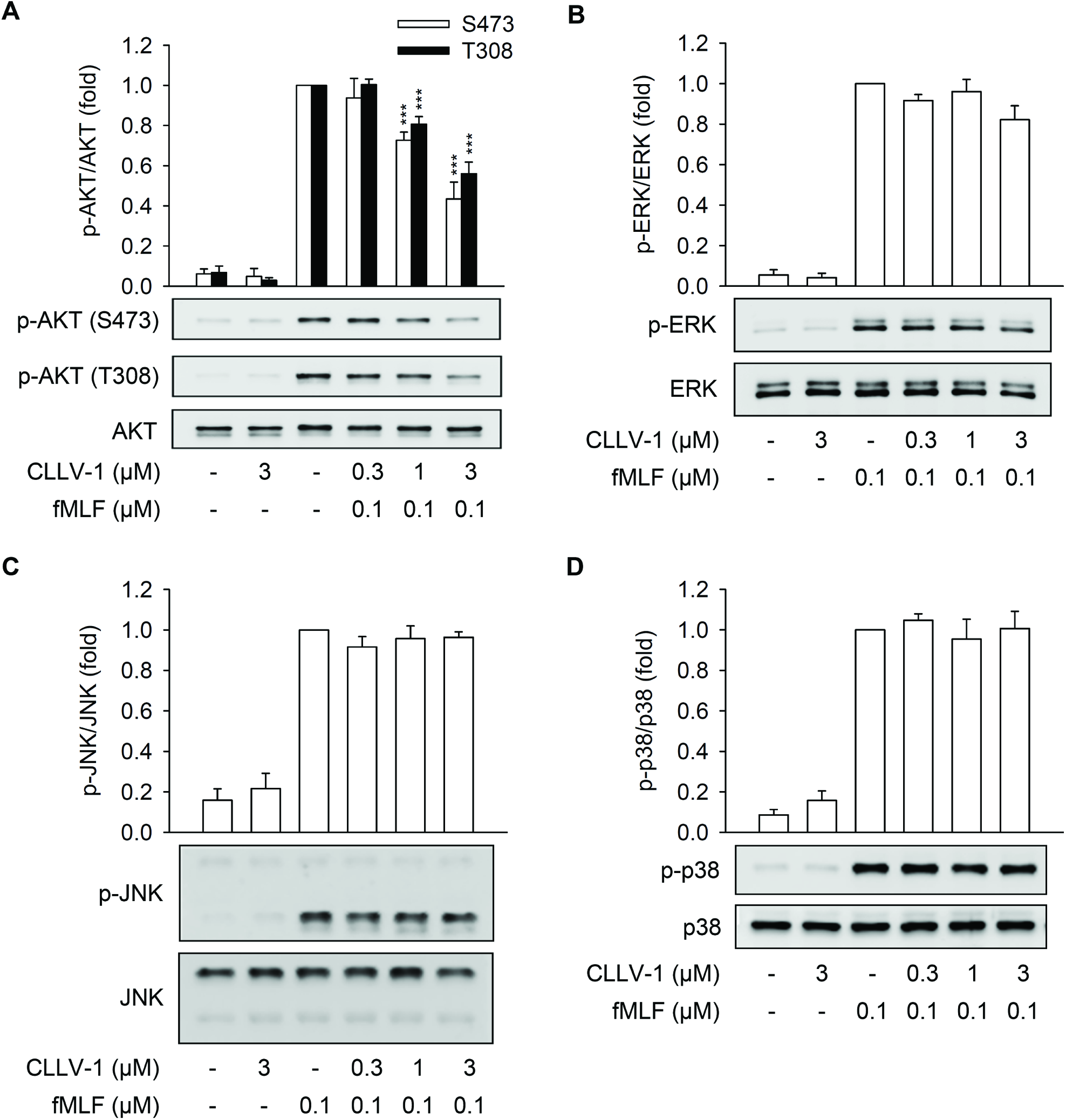
CLLV-1 decreases the phosphorylation of AKT but not MAPKs in fMLF-activated human neutrophils. Human neutrophils were incubated with CLLV-1 (0.3–3 μM) for 5 min before stimulation with or without fMLF (0.1 μM)/CB (1 μg/mL). Phosphorylation of (A) AKT, (B) ERK, (C) JNK, or (D) p38 MAPK was analyzed by immunoblotting using antibodies against the phosphorylated form and the total of each protein. All data are expressed as mean values ± SEM (n = 3); \*\*\**p* < 0.001 compared with the DMSO + fMLF group (Student’s *t*-test).

### CLLV-1 inhibits the inflammatory responses and AKT activation in fMLF-activated dHL-60 cells

HL-60 cells were exposure to DMSO for 5 days to differentiate into neutrophil-like cells (dHL-60 cells). Usage of dHL-60 cells provided enough cells for several biochemical experiments instead of limited primary human neutrophils. The increased FPR1 expression and cellular morphology of dHL-60 were observed to represent the neutrophil-like status (Figure 4A-figure supplement 6). CLLV-1 diminished superoxide anion generation and intracellular ROS formation in fMLF-induced dHL-60 cells. It also repressed the p47^phox^ phosphorylation and F-actin polymerization in fMLF-activated dHL-60 cells (Figure 4B-E-figure supplement 7), suggesting that dHL-60 cells provide a good inflammatory model and that CLLV-1 still restrains the respiratory burst and chemotaxis in fMLF-activated dHL-60 cells. Importantly, CLLV-1 also attenuated fMLF-induced phosphorylation of AKT (Thr308 and Ser473) in dHL-60 (Figure 4F).

**Figure 4.**
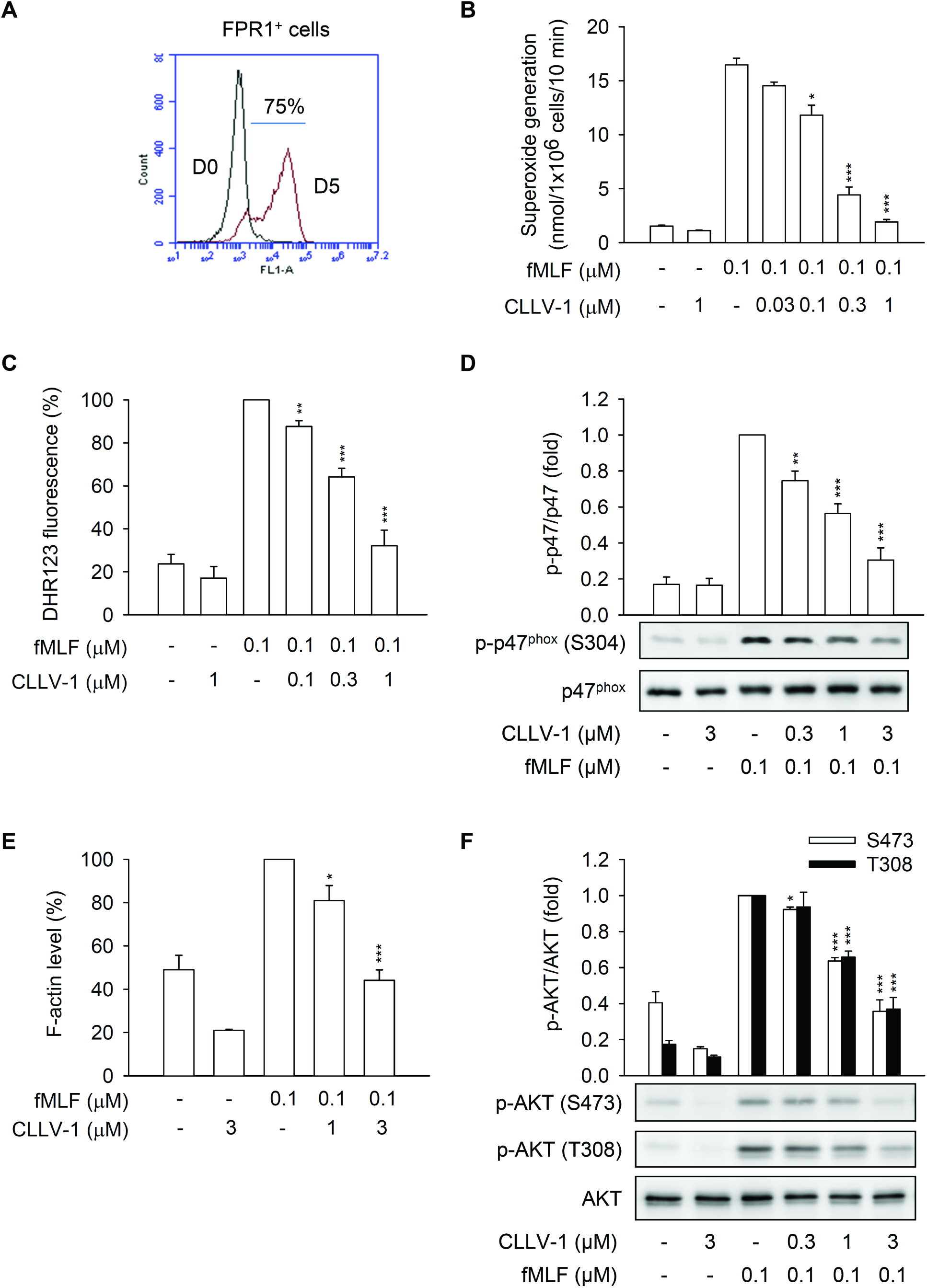
CLLV-1 suppresses fMLF-induced inflammatory responses in differentiated HL-60 (dHL-60) cells. (A) The HL-60 cells were exposed to 1.3% DMSO for 5 days. The differentiation of HL-60 cells by DMSO was examined by flow cytometry, using anti-FPR1 antibodies. (B-F) The dHL-60 cells were preincubated with DMSO or CLLV-1 (0.03–1 μM) and then activated with or without fMLF (0.1 μM)/CB (1 μg/mL). (B) Superoxide anion generation was detected by spectrophotometry at 550 nm, using cytochrome *c* reduction. (C) The intracellular ROS was monitored by flow cytometry, using cell-permeable DHR123. (D) Phosphorylation of p47^phox^ was analyzed by immunoblotting, using antibodies against the phosphorylated (S304) and total p47^phox^. (E) F-actin levels were assayed by flow cytometry, using Alexa Fluor 594 Phalloidin. (F) Phosphorylation of AKT was analyzed by immunoblotting, using antibodies against the phosphorylated (S473 and T308) and total AKT. All data are expressed as mean values ± SEM (n = 3); \**p* < 0.05, \*\**p* < 0.01, and \*\*\**p* < 0.001 compared with the DMSO + fMLF group (Student’s *t*-test).

### CLLV-1 directly alleviates AKT activity

We tested whether the CLLV-1-inhibited AKT phosphorylation is based on altering the activation of upstream kinases of AKT, including phospho-phosphoinositide-dependent protein kinase 1 (p-PDK1), phospho-mammalian target of rapamycin C2 (p-mTORC2), and p-PI3K *(Brazil et al, 2004; Weichhart et al, 2015*). CLLV-1 (0.3-3 μM) failed to affect the phosphorylation of PDK1 (S241), mTORC2 (S2481), and PI3K (Y199 of p85 subunit) in fMLF-activated dHL-60 cells and human neutrophils (Figure 5A-figure supplement 8). The level of PI3K-generated PIP3 was also not changed by CLLV-1 in fMLF-activated dHL-60 cells (Figure 5B-figure supplement 7C). Protein kinase A (PKA) has been shown to attenuate neutrophilic inflammation through inhibiting AKT activation *(Sousa et al, 2010*). However, CLLV-1 did not increase the level of cAMP. The PKA inhibitor, H89, did not reverse the CLLV-1-inhibited superoxide anion generation and elastase release in activated human neutrophils (Figure 5-figure supplement 9).

**Figure 5.**
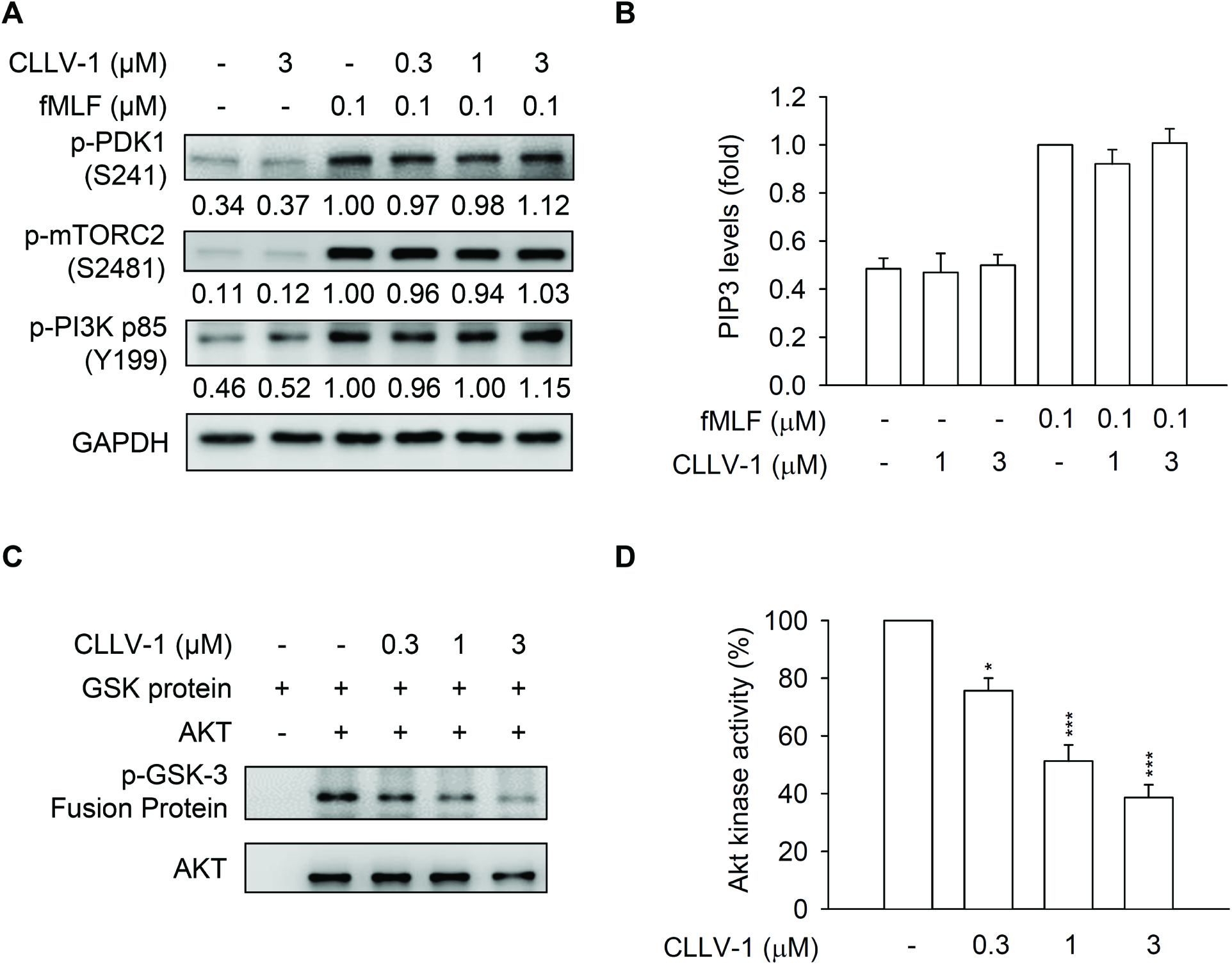
CLLV-1 blocks AKT enzymatic activity but does not affect the AKT upstream kinase. (A-B) dHL-60 cells were preincubated with DMSO or CLLV-1 (0.03–1 μM) and then activated with or without fMLF (0.1 μM)/CB (1 μg/mL). (A) Phosphorylation of AKT upstream kinases, PDK1, mTORC2, and PI3K was determined by immunoblotting, using antibodies against the phosphorylated form and normalized to GAPDH. (B) PIP3 levels were assayed using anti-PIP3 antibodies by flow cytometry. (C) The active AKT proteins were immunoprecipitated using anti-phospho-AKT antibodies and treated with DMSO or CLLV-1 (0.3–3 μM) for 15 min at 30 °C. Subsequently, the GSK-3 fusion protein (AKT substrate) was added for another 30 min. The phospho-GSK-3 fusion protein was examined by immunoblotting. (D) The phosphorylation of the GSK-3 fusion protein was quantified and expressed as a percentage to represent AKT activity. All data are expressed as mean values ± SEM (n = 3); \**p* < 0.05 and \*\*\**p* < 0.001 compared with the DMSO + AKT group (Student’s *t*-test).

We suggest that CLLV-1 may directly target AKT *per se* to repress inflammation in human neutrophils. To explore this hypothesis, the nonradioactive AKT kinase assay was performed *in vitro*. Clearly, our data showed that CLLV-1 (0.3–3 μM) blocked the kinase activity of AKT *in vitro*. MK-2206 was used as a positive control to inhibit AKT activity (Figure 6-figure supplement 5C).

### CLLV-1 covalently reacts with the thiol group of Cys310 in AKT

To determine how CLLV-1 blocked the AKT activity, the molecular docking of CLLV-1 with AKT was performed. Based on CDOCKER and the CHARMm force field, the CLLV-1-AKT binding modes were generated in receptor cavities with 10 poses. The binding of CLLV-1 and AKT with the most favorable energy was estimated with-CDOCKER (−474.532 kcal/mol). The *p*-benzoquinone, aromatic ring, or carboxyl group of CLLV-1 were proposed to interact with the R273, D274, L275, C310, G311, A317, L316, Y315, V320, or V330 residues of AKT (Figure 6A–B). Interestingly, the Cys310 residue of AKT was predicted as a CLLV-1-binding site, and the redox modification of Cys310 in AKT is important for AKT enzymatic activity *(Ahmad et al, 2014; Murata et al, 2003*). Hence, CLLV-1 may bind to the thiol group of Cys310 to interfere with the AKT activity. To address this hypothesis, the reaction between CLLV-1 and synthetic AKT peptides, AKT_304-308_ (ATMKT), AKT_309-313_ (FCGTP), AKT_314-318_ (EYLAP), and AKT_307-328_ (KTFCGTPEYLAPEVLEDNDYGR) were determined by NMR or MS: CLLV-1 covalently reacted with the AKT_309-313,_ exhibiting a new singlet peak at δ 7.00, but did not react with the adjacent AKT_304-308_ and AKT_314-318_, which lacked cysteine residues according to the ^1^H NMR analysis (Figure 6C-figure supplement 10A). The AKT peptide-CLLV-1 adducts were also examined by MS (Figure 6D-E-figure supplement 10B): the molecular masses of AKT_309-313_, AKT_307-328_, and CLLV-1 were 524, 2389, and 268 Da, respectively. If CLLV-1 reacted with the AKT peptides, the proposed molecular masses of the adducts would be 792 Da (AKT_309-313_-CLLV-1) and 2657 Da (AKT_307-328_-CLLV-1). The corresponding signals were observed by MS, including AKT_309-313_ ([M+H]^+^; 524.079), AKT_307-328_ ([M+H]^+^; 2389.124), AKT_309-313_-CLLV-1 ([M_CLL_+H]^+^; 792.244), and AKT_307-328_-CLLV-1 ([M_CLL_+H]^+^; 2655.102). In addition, the sodium adducts (plus 23 Da) were also found, AKT_309-313_-Na ([M+Na]^+^; 546.145), AKT^307-328^-Na ([M+Na]^+^; 2411.258), and AKT_309-313_-CLLV-1-Na ([M_CLL_+Na]^+^; 812.264). Similarly, the results of MS confirmed that CLLV-1 did not react with the adjacent peptides AKT_304-308_ and AKT_314-318_, which do not contain Cys310. These results suggested that CLLV-1 covalently reacted with the thiol group of Cys310 in AKT via an electrophilic addition.

To confirm the effect of CLLV-1 on AKT redox status in cells, we used the alkylation agent, 4-acetamido-4′-maleimidylstilbene-2,2′-disulfonic acid (AMS), to label reduced form of protein along with increasing molecular weight *(Ahmad et al, 2014*). The level of AMS-labeled AKT (reduced form) was increased in fMLF-activated dHL-60 cells. CLLV-1 (0.3–3 μM) dose-dependently decreased the levels of AMS-labeled AKT in fMLF-activated dHL-60 cells (Figure 7A). It has been reported that oxidization of Cys310 residue in AKT diminished its enzymatic activity and increased the interaction between AKT and protein phosphatase 2A (PP2A) *(Ahmad et al, 2014; Durgadoss et al, 2012*). CLLV-1 induced the AKT-PP2A interaction in fMLF-activated dHL-60 cells (Figure 7B). These results suggest that CLLV-1 covalently targets Cys310 of AKT to alleviate AKT activity.

**Figure 6.**
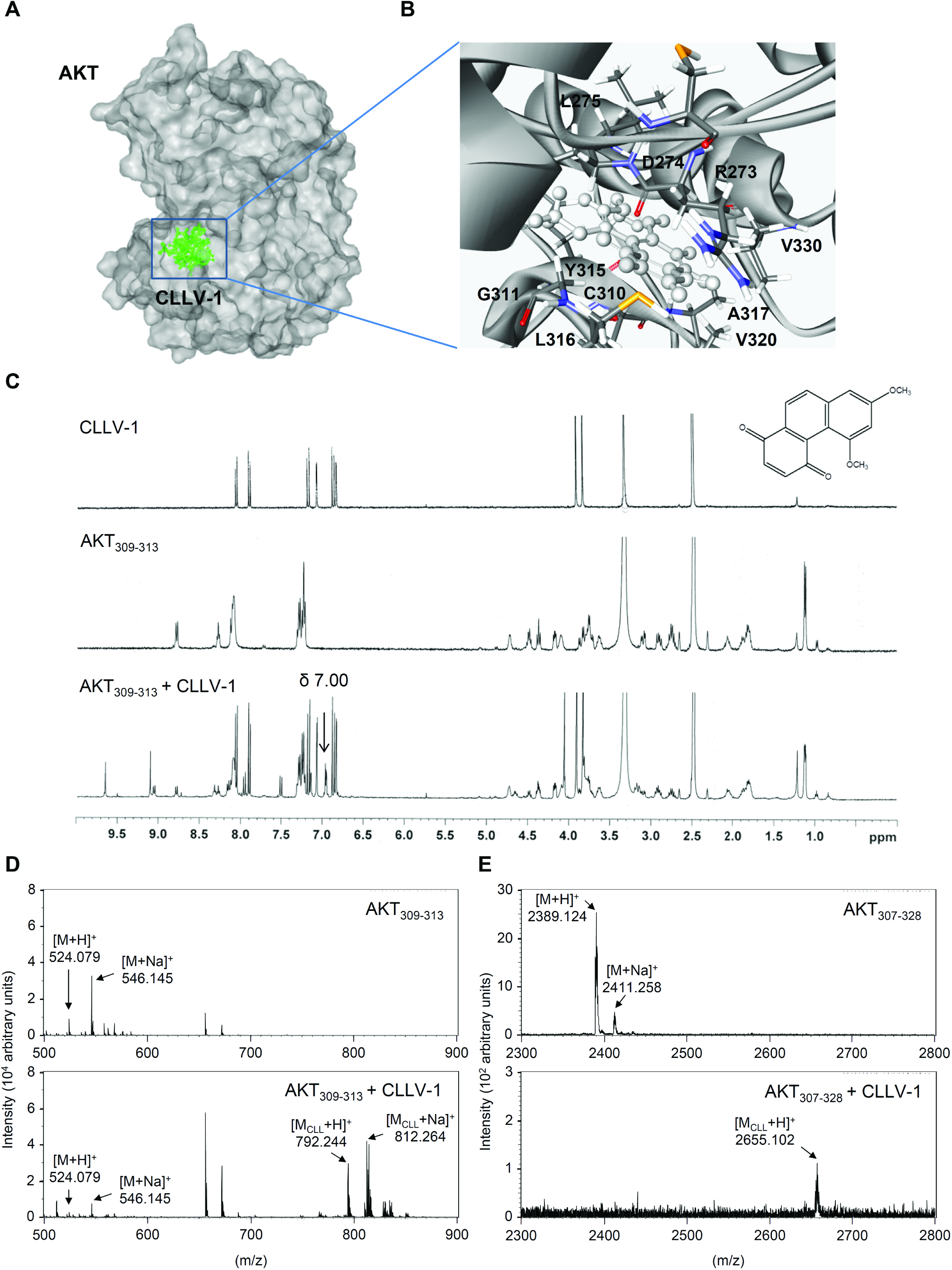
CLLV-1 covalently reacts with the thiol group of an AKT cysteine *in vitro*. (A-B) Docking models of CLLV-1-targeted AKT. Surface presentation demonstrates the structure of AKT (gray). CLLV-1 moieties are colored green and rendered in stick representation (A). Close-up of CLLV-1 docking site (best energy mode) (B). The Figures were prepared using Discovery Studio 4.1. The crystal structure of AKT was downloaded from PDB (accession code 4ekl). The chemical structure of CLLV-1 was drawn by ChemDraw Ultra 9.0. (C) ^1^H NMR spectra of CLLV-1 (upper panel), 	synthetic AKT peptide (AKT_309-313_; FCGTP) (middle panel), and mixtures of CLLV-1 and AKT_309-313_ (lower panel). (D–E) The synthetic AKT peptides AKT_309-313_ and AKT_307-328_ (KTFCGTPEYLAPEVLEDNDYGR) were incubated in the presence or absence of CLLV-1. The molecular mass of the synthetic AKT peptides and their CLLV-1 adducts were detected using MALDI-TOF MS. M, molecular mass of AKT peptides; M_CLL_, molecular mass of adducts of AKT peptides and CLLV-1.

### CLLV-1 attenuates LPS-induced ALI in mice

To examine the anti-inflammatory potential of CLLV-1 *in vivo*, the effects of CLLV-1 on LPS-induced ALI was tested in mice. Intratracheal instillation of LPS showed an increase in pulmonary MPO, a neutrophil infiltration marker, which was inhibited by intraperitoneal injection of CLLV-1 (10 mg/kg) or MK-2206 (10 mg/kg) (Figure 8A). The total protein levels were measured to represent the severity of pulmonary edema. Both CLLV-1 and MK-2206 effectively attenuated the LPS-induced increase of protein levels in the lungs (Figure 8B). The histopathological features of the lungs showed that LPS triggered inflammatory cell infiltration, inter-alveolar septal thickening, and interstitial edema. Moreover, LPS induced the infiltration of cells positive for Ly6G, a specific neutrophil marker, as well as AKT activation in the lungs. Noticeably, administration of CLLV-1 and MK-2206 ameliorated LPS-induced distortion of pulmonary architecture, Ly6G-positive infiltrated neutrophils, and AKT phosphorylation (Figure 8C), suggesting the therapeutic potential of CLLV-1 in neutrophil-dominant lung diseases. Together, CLLV-1 and MK-2206 successfully impeded the inflammatory ALI *in vivo*, supporting that pharmacologically targeting redox modification of AKT is a potential strategy for treating neutrophilic inflammation.

**Figure 7.**
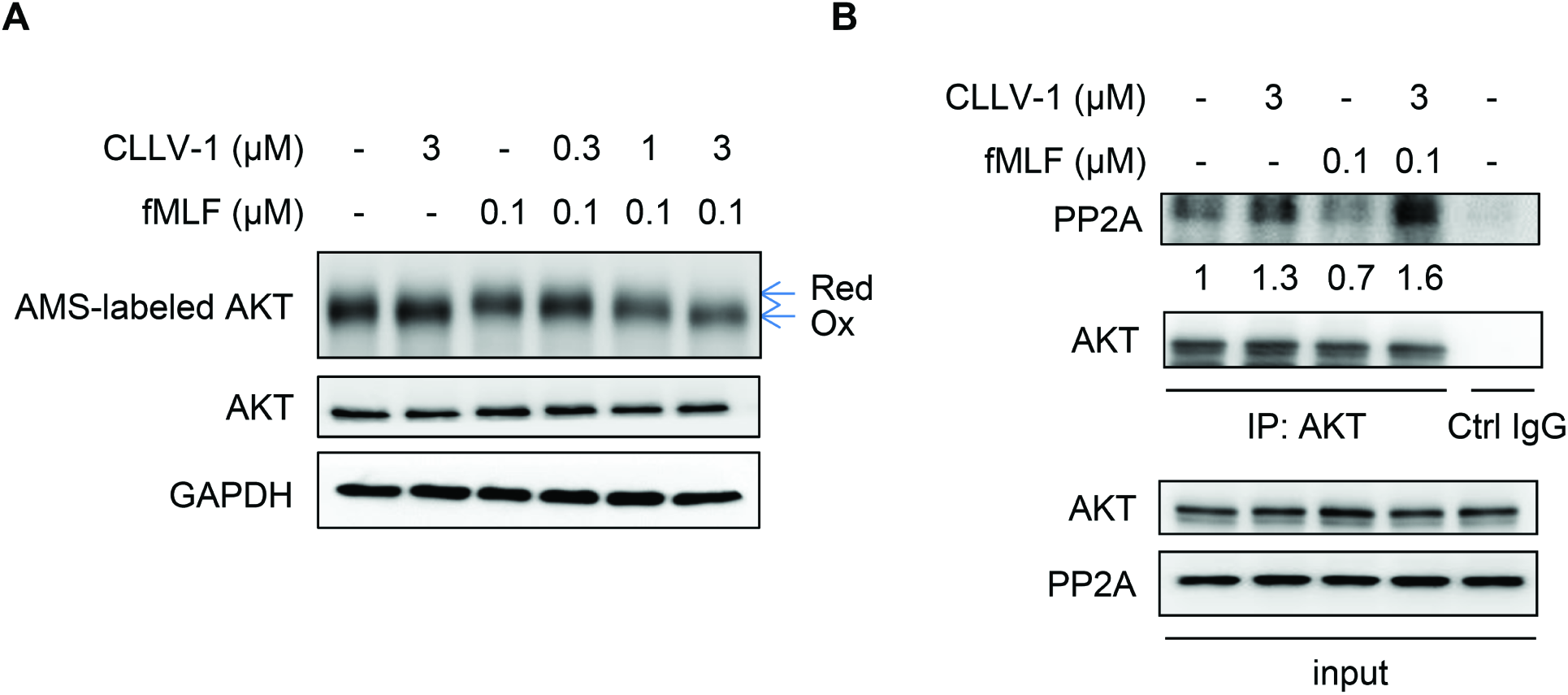
CLLV-1 decreases the AMS-labeled reduced form of AKT and increases the AKT-PP2A interaction in fMLF-activated dHL-60 cells. dHL-60 cells were incubated with CLLV-1 (0.3-3 μM) for 5 min before stimulation with or without fMLF (0.1 μM)/CB (1 μg/mL). (A) The cell lysates were incubated with the thiol-alkylating agent AMS for 12 h on ice and analyzed by Western blotting under reducing (total lysate) or non-reducing conditions (AMS-labeled AKT). The AMS-labeled reduced form of AKT (Red) had a higher molecular weight than oxidized AKT (Oxi). (B) The cell lysates were immunoprecipitated with control (Ctrl) or AKT IgG. The precipitated substances were used for Western blotting of AKT and PP2A.

## Discussion

Neutrophils are the most abundant leukocytes and play a significant role in innate immunity. However, enhanced ROS generation and protease release by activated neutrophils can cause cell and tissue damage *(Nicolas-Avila et al, 2017; Soehnlein et al, 2017*). The AKT pathway is known to be involved in many neutrophil responses, including respiratory burst, degranulation, and chemotaxis; however, the regulatory mechanisms and significance of AKT in neutrophilic inflammation are still elusory *(Chamcheu et al, 2017; Kim et al, 2017; Manning & Toker, 2017; Zhang et al, 2013*). Herein, we show that a synthetic phenanthrenequinone compound, CLLV-1, inhibits inflammatory responses in human neutrophils or neutrophil-like dHL-60 cells triggered by various stimuli. CLLV-1 selectively blocked AKT activity through covalently targeting the Cys310 residue of AKT. Moreover, CLLV-1 attenuated the inflammatory responses in LPS-induced ALI in mice, indicating that it is a potential anti-inflammatory compound and providing an example of disruption of the redox modulation of AKT for treating neutrophil-associated diseases.

AKT is composed of the pleckstrin homology (PH), catalytic kinase, and regulatory domains. With stimulation, AKT is activated and phosphorylated at Thr308 and Ser473 residues, leading to the PH domain becoming more distant from the kinase domain. In contrast, AKT is dephosphorylated and inactivated by PP2A *(Brazil et al, 2004; Weichhart et al, 2015*). Thus far, emerging evidence has supported the important role of redox modification of AKT in conformational dynamics *(Corcoran & Cotter, 2013*). An intramolecular disulfide bond between Cys296 and Cys310 in the catalytic domain of AKT that prompts dephosphorylation by associating with PP2A has been identified; oxidized and dephosphorylated AKT is considered to have lost its kinase activity (*Ahmad et al, 2014; Durgadoss et al, 2012; Murata et al, 2003*). Cys-to-Ser mutation at 296 and 310 in AKT prevents cadmium-inhibited AKT activity and cell survival in neuroblastoma cells (*Ahmad et al, 2014*), supporting the biological significance of the critical Cys in AKT. In the present study, we found that fMLF mitigated the oxidized AKT levels in dHL-60 cells, suggesting that the reduced form of AKT corresponds to its active conformation. CLLV-1 repressed the alkylation agent-labeled AKT levels (reduced form), and the AKT-PP2A interaction was increased by CLLV-1.

CLLV-1 was proposed to interact with Cys310 of AKT by the molecular docking model (Figure 6A). The adduct of AKT_309-313_ peptides and CLLV-1 exhibited a new singlet peak at δ 7.00 in the ^1^H NMR spectrum (Figure 6C). The molecular masses of AKT peptide-CLLV-1 adducts (AKT peptide + CLLV-1 – 2H^+^ Da) were also detected in MALDI-TOF MS, AKT_309-313_-CLLV-1-Na, and AKT_307-328_-CLLV-1 (Figure 6D-E), suggesting that the reaction between AKT and CLLV-1 is an electrophilic addition. It has been reported that thiol-based association between electrophilic compounds and proteins possessed selectivity and specificity. The structural characteristic of proteins and stereochemical structures of electrophiles results in their targeting selectivity (*Dennehy et al, 2006; Mi et al, 2011*). The redox modulations of ERK, JNK, or p38 have been characterized in response to intracellular ROS (*Corcoran & Cotter, 2013*). A low concentration of hydrogen peroxide (< 0.1 μM) induced the oxidation of Cys38 and Cys214 in ERK2 and increased phosphorylation of ERK2; in contrast, doxorubicin-induced oxidation of ERK2 was accompanied by dephosphorylation *(Galli et al, 2008; Luanpitpong et al, 2012*). Upon exposure to a higher concentration of hydrogen peroxide (> 10 μM), the cysteines of JNK2 (Cys41) and p38 (Cys162) were oxidized, followed by activation and phosphorylation (*Galli et al, 2008*). We found that CLLV-1 repressed fMLF-induced AKT, but not ERK, JNK, or p38 activation in human neutrophils (Figure 3), implying its specificity.

Three isoforms of AKT have been identified, AKT1 (PKBα), AKT2 (PKBβ), and AKT3 (PKBγ), sharing 90% identity in kinase domain. Neutrophils contain only AKT1 and AKT2. In AKT isoform-specific knockout mice, overlapping but distinct roles of activated neutrophils have been shown *(Chen et al, 2010; Li et al, 2014*). With fMLF stimulation, AKT2 but not AKT1 translocated to the leading edge in polarized neutrophils and induced superoxide anion generation and p47^phox^ phosphorylation (*Chen et al, 2010*). In the present study, CLLV-1 significantly abrogated AKT activation in response to various stimuli in human neutrophils (Figure 3 and EV2), and the CLLV-1-targeted Cys310 of AKT was identical in AKT1/2/3 (Figure 6), suggesting that CLLV-1 covalently reacts with both AKT1 and AKT2 in neutrophils. It has been reported that AKT controls p47^phox^ phosphorylation and F-actin polymerization to trigger respiratory burst and chemotaxis in neutrophils, respectively *(Chen et al, 2010; El-Benna et al, 2009; Kumar et al, 2014*). CLLV-1 dose-dependently restricted AKT-mediated p47^phox^ phosphorylation and F-actin levels in fMLF-activated human neutrophils and dHL-60 cells (Figures 1, 2, and 4), confirming that CLLV-1-inhibited AKT activity is critical for halting inflammatory activation in human neutrophils.

Developed AKT inhibitors are usually classified as ATP-competitive inhibitors and allosteric inhibitors (*Keane et al, 2014; Nitulescu et al, 2016*). ATP-competitive inhibitors such as GSK690693 significantly suppress AKT activity; however, the off-target effect is still of concern because of the ATP-binding site being highly conserved among members of the AGC kinase family such as PKA and PKC *(Jacinto & Lorberg, 2008*). A growing number of allosteric inhibitors with higher efficacy and specificity, such as MK-2206, is being developed (*Hirai et al, 2010*). The Cys296 and Cys310 residues in the catalytic domain of AKT were identified as allosteric sites for regulating AKT activity (*Lee et al, 2010; Nitulescu et al, 2016; Shearn et al, 2012*). Therefore, the Cys296 and Cys310 residues of AKT are potential therapeutic targets in AKT-associated disorders. In this study, CLLV-1 was found to be a novel allosteric inhibitor of AKT through covalently binding to Cys310 *in vitro* (Figures 5B and 6). This is the first example of restraining neutrophilic inflammation by pharmacologically targeting the redox regulatory site of AKT. CLLV-1 showed anti-inflammatory effects in human neutrophils and ameliorated LPS-primed ALI in mice (Figure 8), supporting the therapeutic potential of CLLV-1 in neutrophilic lung damage. Accordingly, anti-inflammatory drugs that target the redox modification of AKT could potentially be developed.

**Figure 8.**
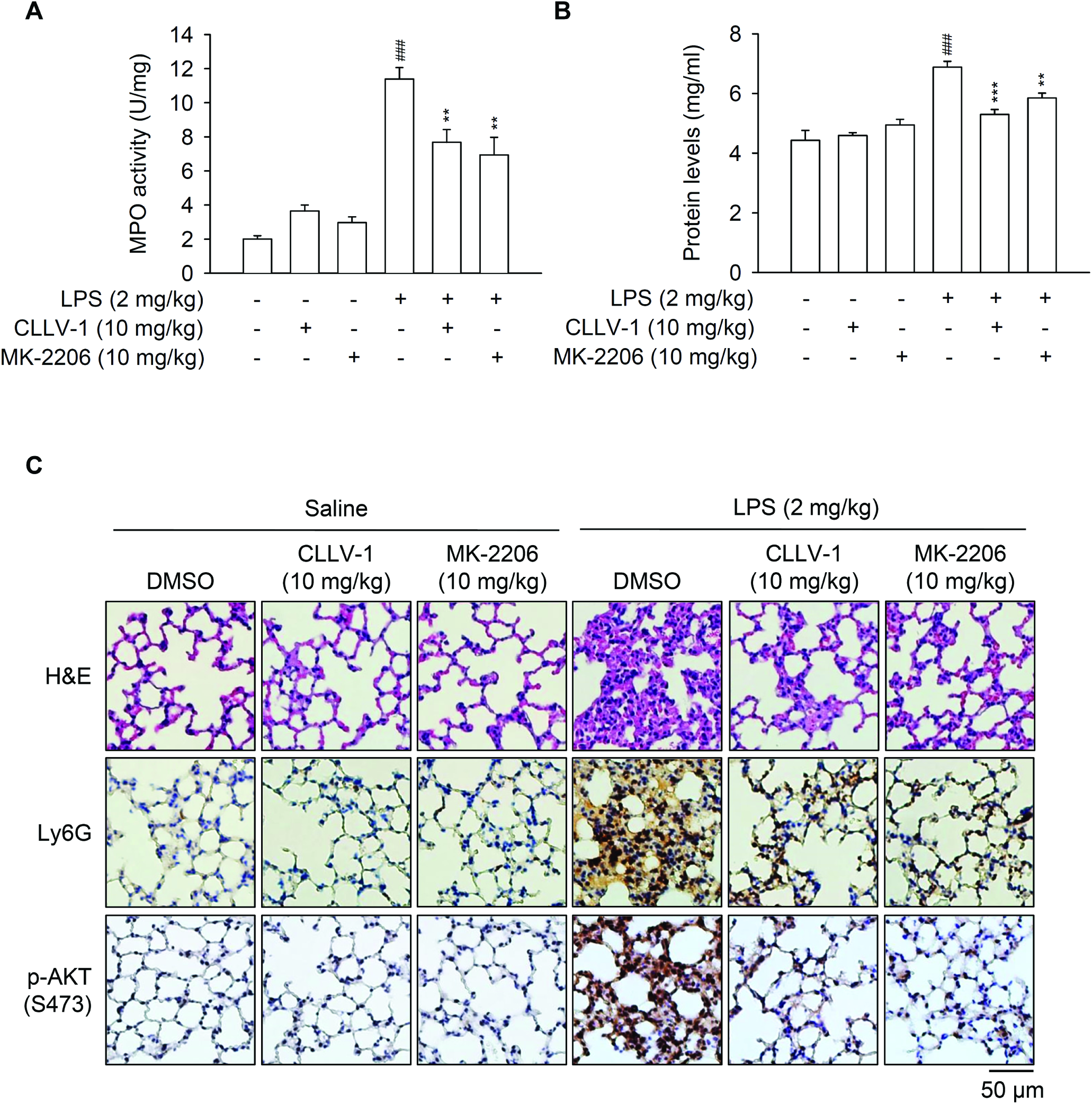
CLLV-1 attenuates LPS-induced ALI in mice. C57BL/6 mice were intraperitoneally injected with CLLV-1 (10 mg/kg), MK-2206 (10 mg/kg), or an equal volume of DMSO for 1 h, and subsequently instilled 2 mg/kg LPS from *E. coli* 0111:B4 or 0.9% saline via tracheostomy. (A-B) Six hours later, lungs were collected and assayed for MPO activity (A) and protein levels (B). (C) Histological examination of lungs. The lung sections were stained with hematoxylin and eosin (H&E), anti-Ly6G antibodies, or anti-pAKT (S473) antibodies by IHC. All data are expressed as mean values ± SEM (n = 6); \*\**p* < 0.01 and \*\*\**p* < 0.001 compared with the LPS + DMSO group; ^###^*p* < 0.001 compared with the DMSO control (Student’s *t*-test).

In summary, we demonstrate that AKT activation plays a critical role in neutrophilic inflammation. Targeting AKT Cys310 using drugs can regulate AKT phosphorylation and activity. Our findings also provide an important example of how a novel derivative of phenanthrenequinone, CLLV-1, restrains neutrophil-associated inflammatory responses by inhibiting AKT activity in a redox-dependent manner (Figure 9).

**Figure 9.**
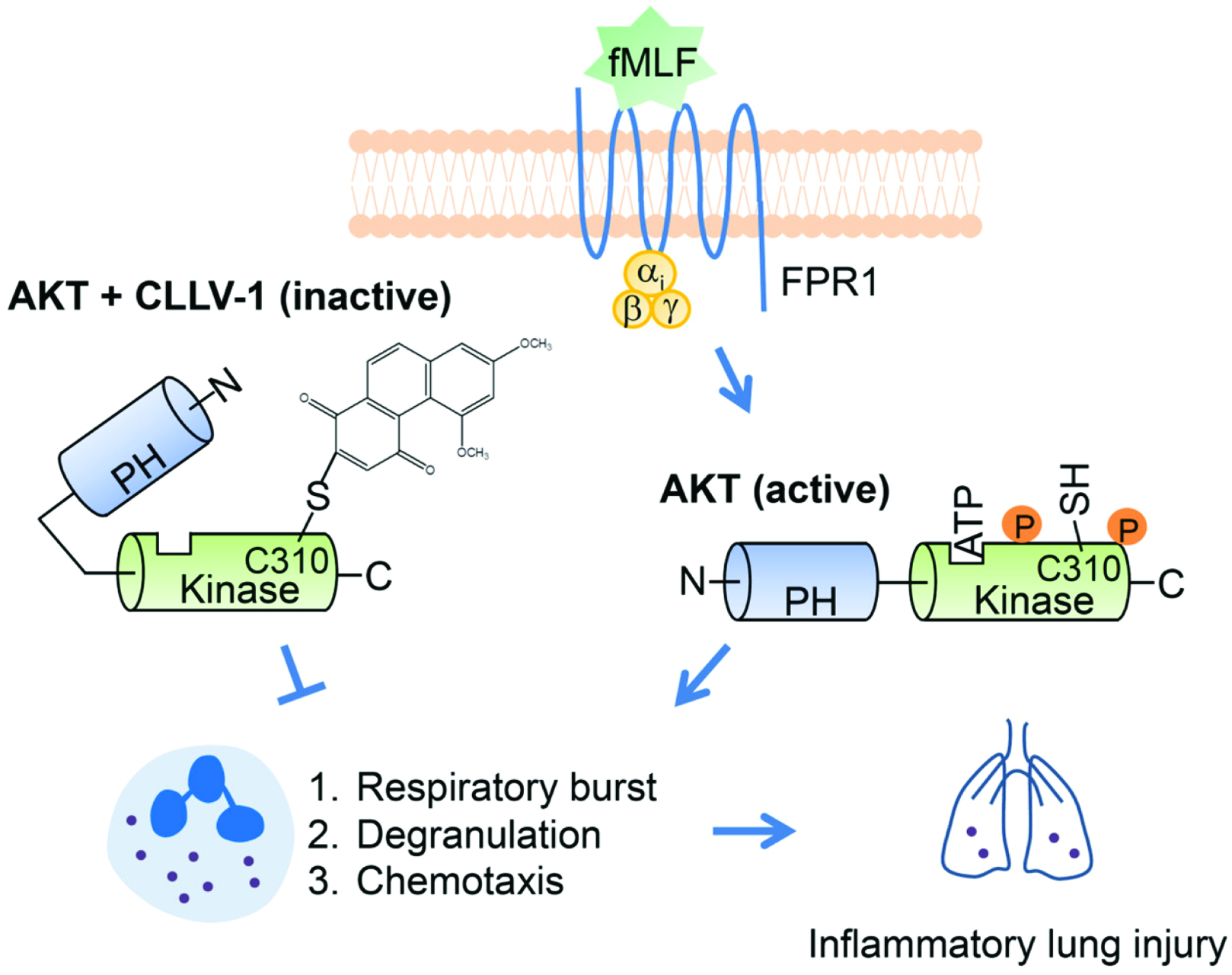
Schematic model of CLLV-1-impeded neutrophilic inflammation. With stimulation, AKT is activated and phosphorylated, leading to the PH domain becoming more distant from the kinase domain. Active AKT is maintained in a reduced manner that contributes to overwhelming inflammatory responses in human neutrophils and neutrophil-dominant inflammatory disorders. Once CLLV-1 is administered, it covalently binds to Cys310 within the kinase domain of AKT, yielding an inactive and less reduced form of AKT. CLLV-1 can inhibit neutrophil activation, including respiratory burst, degranulation, and chemotaxis by modifying the redox state of AKT. Significantly, CLLV-1 ameliorates inflammatory ALI *in vivo*.

## Materials and methods

### Reagents

CLLV-1 was synthesized by our group (*Lee et al, 2008*). The CLLV-1 structure was determined by ^1^H nuclear magnetic resonance (NMR) spectrum analysis (Figure 6C). The purity of CLLV-1 was higher than 96% as determined by HPLC. MK-2206 was purchased from Selleckchem (Houston, TX, USA). WKYMVm was purchased from Tocris Bioscience (Ellisville, MO, USA). FITC-labeled anti-CD11b, anti-Ly-6G, and anti-MPO antibodies were purchased from eBioscience (San Diego, CA, USA). The antibodies against p38 or p47^phox^ and protein G beads were purchased from Santa Cruz Biotechnology (Santa Cruz, CA, USA). Anti-p-p47^phox^ antibodies were purchased from Abcam (Cambridge, MA, USA). Anti-PIP3 antibody was purchased from Echelon Biosciences (Salt Lake City, UT, USA). Other antibodies and the nonradioactive AKT kinase assay kit were purchased from Cell Signaling (Beverly, MA, USA). RPMI 1640, DMEM, L-glutamine, Antibiotic-Antimycotic, DHR123, fNLFNYK, Alexa Fluor 594 Phalloidin, and Hoechst 33342 were purchased from Thermo Fisher Scientific (Waltham, MA, USA). Fetal bovine serum (FBS) was purchased from Biological Industries (Beth Haemek, Israel). Other regents were purchased from Sigma-Aldrich (St. Louis, MO, USA).

### Neutrophil isolation and cell culture

The procedure of neutrophil isolation was approved by the Institutional Review Board at Chang Gung Memorial Hospital. Neutrophils were isolated by dextran sedimentation and Ficoll-Hypaque centrifugation. Blood was obtained from healthy volunteers (20–35 years old), and written informed consent was obtained from every volunteer. The purified neutrophils contained > 98% viable cells, determined by trypan blue exclusion assay (*Chen et al, 2016*).

Endothelial cells (bEnd.3) were obtained from the Bioresource Collection and Research Centre (Hsinchu, Taiwan) and cultured in DMEM media supplemented with 10% FBS and 1× Antibiotic-Antimycotic. The human promyelocytic leukemic HL-60 cell line was purchased from ATCC and cultured in RPMI 1640 medium supplemented with 20% FBS, 2 mM L-glutamine, and 1× Antibiotic-Antimycotic. Both cell lines were grown in a humidified atmosphere (37 °C, 5% CO_2_). The HL-60 cells were differentiated to neutrophil-like cells (dHL-60 cells) by a 5-day treatment with 1.3% DMSO in the growth medium.

### Extracellular superoxide anion production

Human neutrophils (6 × 10^5^ cells/mL) or dHL-60 cells (1 × 10^6^ cells/mL) were equilibrated with 0.5 mg/mL ferricytochrome *c* and 1 mM Ca^2+^ at 37 °C for 5 min. The cells were then preincubated with DMSO, CLLV-1, or MK-2206 and stimulated with fMLF, NaF, or WKYMVm before being primed with cytochalasin B (CB; 1 or 2 μg/ml) for 3 min. The superoxide anion was determined using a spectrophotometer (Hitachi, Tokyo, Japan) at 550 nm.

### Intracellular ROS formation

DHR123 (2 μM)-labeled human neutrophils or dHL-60 cells (1 × 10^6^ cells/mL) were incubated at 37 °C for 10 min. Subsequently, the cells were pretreated with DMSO or CLLV-1 for 5 min and activated by fMLF (0.1 μM)/CB (1 μg/mL) for another 5 min. The intracellular ROS formation was determined using a flow cytometer (BD Bioscience).

### Elastase release

Human neutrophils (6 × 10^5^ cells/mL) were equilibrated with an elastase substrate (MeO-Suc-Ala-Ala-Pro-Val-p-nitroanilide; 100 μM) at 37 °C for 5 min and then incubated with DMSO, CLLV-1, or MK-2206 for 5 min. The cells were then activated by fMLF, NaF, WKYMVm, IL-8, or LTB_4_ for a further 10 min. CB (0.5 or 2 μg/mL) was added 3 min before stimulation. Elastase release was determined by spectrophotometry at 405 nm.

### CD11b expression

Neutrophils (5 × 10^6^ cells/mL) were preincubated with DMSO or CLLV-1 for 5 min and activated by fMLF (0.1 μM)/CB (0.5 μg/mL) for another 5 min. The cell pellets were then resuspended in 5% bovine serum albumin (BSA) containing FITC-labeled anti-CD11b antibodies (1 μg) at 4 °C for 90 min. The fluorescence intensity was measured by flow cytometry.

### Superoxide anion-scavenging assay

The superoxide anion-scavenging effect of CLLV-1 was examined in a cell-free xanthine/xanthine oxidase system. The assay buffer contained 50 mM Tris (pH 7.4), 0.3 mM WST-1, and 0.02 U/mL xanthine oxidase. WST-1 was reduced by the superoxide anion after adding 0.1 mM xanthine to the assay buffer at 30 °C. The absorbance was measured at 450 nm.

### NADPH oxidase activity

Neutrophils were mixed with 1 mM phenylmethylsulphonyl fluoride for 30 min at 4 °C and sonicated in relaxation buffer (10 mM piperazine-N,N′-bis(2-ethanesulfonic acid) PIPES, 100 mM KCl, 3 mM NaCl, 3.5 mM MgCl_2_, 1 mM ethylene glycol-bis(β-aminoethyl ether)-N,N,N′,N′-tetraacetic acid (EGTA), and 1 mM ATP; pH 7.3) to produce cytosolic and plasma membrane fractions. NADPH oxidase was activated by the addition of 100 μM SDS before incubation with DMSO or CLLV-1 for 2 min as described.

### Elastase activity

Human neutrophils (6 × 10^5^ cells/mL) were activated by the addition of fMLF (0.1 μM) in the presence of CB (2.5 μg/mL) for 15 min at 37 °C. Elastase was obtained from the supernatant of the cells after they were centrifuged at 1000 *g* for 5 min at 4 °C. Then, the supernatant was equilibrated at 37 °C for 2 min and incubated with or without CLLV-1 for 5 min. After incubation, elastase substrate (100 μM) was added to the reaction solutions. The changes in absorbance were continuously monitored for 10 min at 405 nm to determine the elastase activity.

### LDH release

The CytoTox 96 non-radioactive cytotoxicity assay (Promega, Maddison, WI, USA) was used to determine the LDH level. Human neutrophils were treated with CLLV-1 (1 or 3 μM) for 15 or 60 min. The supernatant was obtained for the LDH assay. LDH release was expressed as a percentage of the amount of enzyme liberated following incubation of human neutrophils with 0.1% Triton X-100 for 30 min at 37 °C.

### Receptor-binding assay

Receptor binding was assayed by FACScan analysis of the binding of fNLFNYK, a fluorescent analog of fMLF. Cells were incubated with CLLV-1 for 5 min at 4 °C and labeled with fNLFNYK. After 30 min, the cells were pelleted, resuspended in ice-cold HBSS, and immediately analyzed using a flow cytometer (FACSCalibur™; BD Bioscience).

### Western blotting

Human neutrophils were preincubated with DMSO or CLLV-1 at 37 °C for 5 min and then activated by fMLF, NaF, WKYMVm, IL-8, or LTB_4_ before being primed with CB. The reaction was stopped using the sample buffer (62.5 mM pH 6.8 Tris-HCl, 4% SDS, 5% β-mercaptoethanol, 2.5 mM Na_3_VO_4_, 0.00125% bromophenol blue, 10 mM di-N-pentyl phthalate, and 8.75% glycerol) at 100 °C for 15 min. The cell lysates were separated by SDS-polyacrylamide gel electrophoresis (PAGE), and assayed by immunoblotting with the corresponding antibodies, followed by incubation with horseradish peroxidase-conjugated secondary anti-rabbit or anti-mouse antibodies. The labeled proteins were measured using an enhanced chemiluminescence system (Amersham Biosciences, Piscataway, NJ, USA).

### PIP3 and F-actin expression

Neutrophils or dHL-60 cells (5 × 10^6^ cells/mL) were preincubated with DMSO or CLLV-1 for 5 min and then activated by fMLF (0.1 μM)/CB (1 μg/mL). The reaction was stopped by 4% paraformaldehyde at 25 °C for 20 min and then permeabilized with 0.1% Triton-X-100. For F-actin staining, cells were incubated with Alexa Fluor 594 Phalloidin in Hank’s balanced salt solution (HBSS) containing 2% BSA at 25 °C for 60 min. For PIP3 expression, cells were incubated with anti-PIP3 antibodies and FITC-labeled anti-mouse IgG antibodies in HBSS containing 2% BSA at 25 °C for 60 min, respectively. The fluorescence intensity was monitored using flow cytometry.

### AKT kinase assay

The AKT activity was determined using the non-radioactive AKT kinase assay kit according to the manufacturer’s protocol. In brief, dHL-60 cells were activated by fMLF (0.1 μM)/CB (1 μg/mL), and the active AKT in the cell lysate was immunoprecipitated with immobilized AKT primary antibodies. The precipitated AKT was treated with DMSO, CLLV-1, or MK-2206 at 30 °C for 15 min and then incubated with ATP and GSK-3 fusion protein as a kinase substrate at 30 °C for 30 min. The reaction was stopped by 3× sodium dodecyl sulfate (SDS) sample buffer at 100 °C for 5 min. The phosphorylation of the GSK-3 fusion protein was determined by western blot.

### cAMP concentration

The concentration of cAMP was determined with an enzyme immunoassay kit (GE Healthcare, Little Chalfont, UK). Neutrophils were treated with DMSO or CLLV-1 at 37 °C for 5 min and then activated by 0.1 μM fMLF for another 1 min. The reaction was terminated by adding 0.5% dodecyltrimethylammonium bromide. After samples were centrifuged at 3000 *g* for 5 min at 4 °C, the supernatants were analyzed for cAMP according to the manufacturer’s instructions.

### Molecular docking

CLLV-1 was docked on AKT proteins by docking optimization (CDOCKER) and optimized with the CHARMm force field using Discovery Studio 4.1 (DS) (BIOVIA, San Diego, CA). The binding of CLLV-1 and AKT1 with the most favorable energy was estimated with-CDOCKER (-kcal/mol). The crystal structure of AKT1 was obtained from the Protein Data Bank (PDB; accession code 4ekl). The 3D structure of CLLV-1 was drawn using ChemDraw Ultra 9.0.

### NMR spectrum analysis

CLLV-1 (1 mg) or synthetic AKT peptides (1 mg) were dissolved in 0.5 mL DMSO-*d6*. The mixtures of CLLV-1 (0.5 mg) and AKT peptides (1 mg) were vigorously mixed in 0.6 mL DMSO-*d6* and incubated at 25 °C for 1 h. The ^1^H NMR spectra were acquired using a Bruker AVANCE-400MHz FT-NMR spectrometer (Bruker BioSpin GmbH, Billerica, MA).

### MS analysis

Synthetic AKT peptides were dissolved in PBS. The mixtures of AKT peptides (120 μM) and CLLV-1 (60 μM) were incubated at 25 °C for 2 h. The AKT peptides and their CLLV-1 adducts were detected using matrix-assisted laser desorption/ionization time of flight mass spectrometer (MALDI-TOF MS). The AKT peptides and their CLLV-1 adducts were mixed with α-Cyano-4-hydroxycinnamic acid (CHCA) matrix (2 mg/mL in 80% acetonitrile containing 0.1% trichloroacetic acid) and loaded onto an MTP AnchorChip™ 600/384 TF (Bruker Daltonics GmbH, Bremen, Germany). After the crystallization of the peptides and the matrix, the samples were analyzed by an Ultraflex™ MALDI-TOF MS (Bruker Daltonics GmbH), controlled by the FlexControl software (v.2.2; Bruker Daltonics GmbH). Data processing was performed and monoisotopic peptide mass was acquired using the FlexAnalysis 2.4 peak-picking software (Bruker Daltonics GmbH).

### AMS labeling assay

The redox states of the proteins were examined by conjugating free thiol with AMS (*Ahmad et al, 2014*). The cells were lysed in the buffer (50 mM Tris, pH 7.4, 150 mM NaCl, 0.5% Triton-X-100, and 1× protease inhibitor cocktail) and centrifuged at 12,000 *g* for 10 min. The supernatants were incubated with 30 mM AMS at 4 °C for 24 h and then mixed with non-reducing sample buffer (62.5 mM pH 6.8 Tris-HCl, 4% SDS, 0.00125% bromophenol blue, and 8.75% glycerol) at 37 °C for 10 min. The redox states of the proteins were determined by immunoblotting.

### Immunoprecipitation

Cells were lysed in the buffer (50 mM Tris, pH 7.4, 150 mM NaCl, 0.5% Triton X-100, 1X protease inhibitor cocktail) and centrifuged at 12,000 *g* for 10 min. The supernatants were incubated with AKT antibodies bound to protein G beads. The beads were washed with buffer and the precipitated proteins were assayed by immunoblotting.

### Neutrophil adhesion and chemotactic migration assays

The bEnd.3 ECs were activated with LPS (2 μg/mL) for 4 h. Hoechst 33342-labeled neutrophils were preincubated with DMSO or CLLV-1 for 5 min and activated by fMLF (0.1 μM)/CB (1 μg/mL) for another 5 min. Activated neutrophils were then co-cultured with LPS-pre-activated bEnd.3 ECs for 30 min. After gently washing, neutrophils adhering to bEnd.3 ECs were randomly counted in 4 fields by microscopy (IX81; Olympus, Center Valley, PA, USA) (*Chen et al, 2016*).

DMSO- or CLLV-1-pretreated neutrophils in the top microchemotaxis chamber (Merck Millipore, Darmstadt, Germany) were placed into the bottom well containing 0.1 μM fMLF. After 90 min, the migrated neutrophils were counted.

### LPS-induced ALI

Seven to eight weeks old C57BL/6 male mice were intraperitoneally injected with CLLV-1 (10 mg/kg), MK-2206 (10 mg/kg) or an equal volume of DMSO. After 1 h, tracheostomy was performed under anesthesia. Mice were instilled with 2 mg/kg LPS (*Escherichia coli* 0111:B4) or 0.9% saline. After 6 h, the lungs were fixed in 10% formalin for immunohistochemistry or frozen for myeloperoxidase (MPO) activity.

### MPO activity

The lung tissues were immersed in 10 mM PBS, pH 6.0, with 0.5% hexadecyltrimethylammonium bromide and sonicated by a homogenizer. The MPO activity was determined using MPO substrate buffer (PBS, pH 6.0, 0.2 mg/mL o-dianisidine hydrochloride, and 0.001% hydrogen peroxide) and monitored the absorbance at 405 nm by a spectrophotometer. The serial concentration of human MPO was used as a standard curve to calculate the MPO activity. Total protein levels were measured by the Bradford protein assay (Bio-Rad, Hercules, CA, USA). The final MPO activity was normalized to the corresponding protein concentration (U/mg).

### Immunohistochemistry (IHC)

Formalin-fixed paraffin-embedded tissue sections were used for IHC. All slides were stained with hematoxylin and eosin (H&E) for morphologic determination. The anti-MPO antibodies, anti-Ly6G antibodies, anti-pAKT (S473) antibodies, or the SuperPicture kit (Thermo Fisher Scientific) were used as the primary or secondary antibodies, following a previously published protocol by Pathology Core of Chang Gung University (*Yuan et al, 2006*).

### Statistical analysis

Data are presented as means ± SEM. Statistical analysis was performed with SigmaPlot (Systat Software) using Student’s *t*-test. *P* < 0.05 was considered statistically significant.

## Contributors

P.-J.C. designed and performed most experiments. I.-L. K., C.-L. L., F.-R. C., Y.-C. Wu., Y.-L. L., C.-C. W., Y.-F. T., C.-Y. Lin. and C.-Y. P. helped to perform experiments and analyzed the data. C.-L. L., H.-C. H., F.-R. C. and Y.-C. W. isolated and provide CLLV-1. P.-J.C. and T.-L.H. wrote and completed the manuscript. T.-L.H. supervised the entire study.

## Figure supplement legends

**Figure supplement 1.** CLLV-1 has no effect on cell viability, superoxide anion scavenging, NADPH oxidase activity, elastase activity, and FPR1 binding. (A) Neutrophils were incubated with DMSO or CLLV-1 (1 or 3 μM) for 15 and 60 min. Cytotoxicity was evaluated by LDH release as a percentage of total LDH release. (B) The superoxide anion scavenging effect of CLLV-1 (0.1–3 μM) was assayed in a cell-free xanthine/xanthine oxidase system. Reduction of WST-1 was measured spectrophotometrically at 450 nm. SOD (0.5 or 20 U/mL) was used as the positive control. (C) The subcellular NADPH oxidase activity was investigated in an SDS-reconstituted system. A reaction mixture of neutrophil cytosolic fraction and membrane fraction at 30 °C was activated by the addition of 100 μM SDS before incubation with DMSO, CLLV-1 (0.3 or 3 μM), or DPI (10 μM) for 2 min. The reaction was initiated by adding 0.16 mM NADPH, and the changes in absorbance at 550 nm caused by the addition of 0.16 mM NADPH were measured. (D) Human neutrophils were pretreated with fMLF/CB for 15 min. The supernatant was obtained and incubated with DMSO or CLLV-1 (0.1–3 μM) before the addition of substrate (100 μM). Elastase activity was measured spectrophotometrically at 405 nm. (E) Neutrophils were preincubated with DMSO, CLLV-1 (0.1–3 μM), or fMLF (10 μM) for 5 min before labeling with the fluorescence-labeled peptide fNLFNYK (4 nM) for another 30 min. The fluorescence was determined by flow cytometry. All data are expressed as mean values ± SEM (n = 3); \*\*\**p* < 0.001 compared with the control (Student’s *t*-test).

**Figure supplement 2.** The representative histograms show an inhibitory effect of CLLV-1 on human neutrophils adhering to ECs. Hoechst 33342-labeled neutrophils were incubated with DMSO or CLLV-1 (0.1–3 μM) for 5 min before activation with fMLF (0.1 μM)/CB (1 μg/mL). Activated neutrophils were then co-cultured with LPS-pre-activated ECs for 30 min. Neutrophils adhering to ECs were detected using microscopy.

**Figure supplement 3.** CLLV-1 represses superoxide anion generation or elastase release in NaF-, WKYMVm-, IL-8-, or LTB_4_-activated human neutrophils. Human neutrophils were preincubated with DMSO or CLLV-1 (0.03–3 μM) and then activated with or without (A,C) NaF (20 mM)/CB (2 μg/mL), (B,D) WKYMVm (1 nM)/CB (1 μg/mL), (E) IL-8 (50 ng/mL)/CB (2 μg/mL), or (F) LTB_4_ (0.1 μM)/CB (0.5 μg/mL). Superoxide anion generation (A–B) and elastase release (C–F) were assayed using cytochrome *c* reduction and elastase substrate by spectrophotometry at 550 nm and 405 nm, respectively. All data are expressed as mean values ± SEM (n = 3); \*\**p* < 0.01 and \*\*\**p* < 0.001 compared with the DMSO + fMLF group (Student’s *t*-test).

**Figure supplement 4.** CLLV-1 inhibits the phosphorylation of AKT in NaF-, WKYMVm-, IL-8-, or LTB_4_-activated human neutrophils. Human neutrophils were preincubated with DMSO or CLLV-1 (0.03–3 μM) and then activated with or without (A) NaF (20 mM)/CB (2 μg/mL), (B) WKYMVm (1 nM)/CB (1 μg/mL), (C) IL-8 (50 ng/mL)/CB (2 μg/mL), or (D) LTB_4_ (0.1 μM)/CB (0.5 μg/mL). Phosphorylation of AKT was analyzed by immunoblotting, using antibodies against phosphorylated (S473 and T308) and total AKT. All data are expressed as mean values ± SEM (n = 3); \**p* < 0.05, \*\**p* < 0.01, and \*\*\**p* < 0.001 compared with the DMSO + fMLF group (Student’s *t*-test).

**Figure supplement 5.** MK-2206 suppresses superoxide anion generation and elastase release in fMLF-activated human neutrophils and AKT activity *in vitro*. Human neutrophils were preincubated with DMSO or MK-2206 (0.3-10 μM) and then activated with or without fMLF (0.1 μM)/CB (1 μg/mL). (A) Superoxide anion generation was detected using cytochrome *c* reduction by spectrophotometry at 550 nm. (B) Elastase release was measured at 405 nm. (C) The active AKT proteins were immunoprecipitated with phospho-AKT antibodies and treated with DMSO, CLLV-1 (1 or 3 μM), or MK-2206 (0.3–3 μM) for 15 min at 30 °C, followed by treatment with GSK-3 fusion protein (AKT substrate) for another 30 min. The phospho-GSK-3 fusion protein was examined by immunoblotting. All data are expressed as mean values ± SEM (n = 3); \**p* < 0.05, \*\**p* < 0.01, and \*\*\**p* < 0.001 compared with the DMSO + fMLF group (A or B) or GSK protein + AKT group (C) (Student’s *t*-test).

**Figure supplement 6.** The cellular morphology of differentiated HL-60 (dHL-60) cells. The cellular morphologies of HL-60 cells or DMSO-differentiated dHL-60 cells were observed by microscopy for 5 days (D5).

**Figure supplement 7.** The representative histograms of flow cytometric analysis show an inhibitory effect of CLLV-1 on ROS formation and F-actin assembly, but not on PIP3 expression in fMLF-activated dHL-60 cells. (A) DHR123 was used to detect the intracellular ROS. (B) Alexa Fluor 594 Phalloidin was used to examine the F-actin assembly. (C) Anti-PIP3 antibodies and FITC-labeled anti-mouse IgG antibodies were used to measure PIP3 expression.

**Figure supplement 8.** CLLV-1 does not affect the AKT upstream kinases in fMLF-activated human neutrophils. Phosphorylation of AKT upstream kinases, PDK1, mTORC2, and PI3K was determined by immunoblotting using antibodies against the phosphorylated form and normalized to GAPDH.

**Figure supplement 9.** The anti-inflammatory effects of CLLV-1 are independent of cAMP levels and PKA in fMLF-activated human neutrophils. (A-B) Human neutrophils were preincubated for 5 min with H_2_O or H89 (3 μM) before the addition of DMSO, CLLV-1 (0.3 μM), or PGE1 (3 μM) and then activated by fMLF (0.1 μM)/CB (1 μg/mL). (A) Superoxide anion generation was detected using cytochrome *c* reduction by spectrophotometry at 550 nm. (B) Elastase release was measured using elastase substrate by spectrophotometry at 405 nm. (C) Human neutrophils were incubated with DMSO or CLLV-1 (0.1 or 1 μM) for 5 min before stimulation with or without fMLF (0.1 μM)/CB (1 μg/mL) for another 1 min. The cAMP levels were measured by enzyme immunoassay kits. All data are expressed as mean values ± SEM (n = 3); \*\*\**p* < 0.001 compared with the H_2_O group (Student’s *t*-test).

**Figure supplement 10.** The chemical interaction of CLLV-1 and synthetic AKT peptides. (A) ^1^H NMR spectra of CLLV-1, synthetic AKT peptides (304-308 or 314-318), and mixtures of CLLV-1 and AKT peptides. (B) The synthetic AKT peptides AKT_304-308_ or AKT_314-318_ were incubated in the presence or absence of CLLV-1. The molecular masses of the synthetic AKT peptides and their CLLV-1 adducts were detected using MALDI-TOF MS. M, molecular mass of AKT peptides.

